# Decoupling Detection and Classification to Improve Morphological Phenotype Analysis of Sickle Red Blood Cells in Full-Scope Microscopy

**DOI:** 10.64898/2026.03.31.715578

**Authors:** Suqiang Ma, Mengjia Xu, Ming Dao, He Li

## Abstract

Microscopy-based analysis of red blood cell (RBC) morphology is widely used to study phenotypes in sickle cell disease (SCD). Although AI models have been developed to automate classification, most are trained on pre-cropped single-cell images and thus struggle with full-scope microscopic images containing densely packed cells and diverse morphologies, which require both accurate detection and fine-grained classification. We propose an end-to-end computational framework to identify individual RBCs in full-scope microscopy images and classify them into five morphological categories: discocytes (DO), echinocytes (E), elongated and sickle-shaped cells (ES), granular cells (G), and reticulocytes (R). We first evaluate advanced detection-classification models, including You Only Look Once (YOLO) and Detection Transformers (DETR), and demonstrate that while these models effectively detect cells, their classification performance falls short of specialized classifiers trained on single-cell images, particularly for minority phenotypes. To address this limitation, we introduce a two-step framework in which a YOLO-based detector localizes and crops individual cells from full-scope images, followed by a fine-tuned DenseNet121 ensemble classifier that assigns each cell to one of the five morphological categories. The proposed framework achieves a detection-level F1-score of 0.9661 and a weighted-average classification F1-score of 0.9708, with an overall classification accuracy of 97.06%. Compared with the single-step YOLO26n baseline, the two-step pipeline yields a macro-average F1-score improvement of +0.1675, with particularly substantial gains for minority classes (E: +0.1623; G: +0.2774; R: +0.2603). Overall, this hybrid framework demonstrates a practical strategy for adapting fast, general-purpose detection models to domain-specific biomedical tasks by combining them with specialized classifiers, delivering both efficiency and high accuracy for scientific and clinical image analysis.

## 1 Introduction

Biomedical imaging has emerged as a crucial tool for modern biological and clinical research, significantly enhancing our ability to visualize, analyze, and interpret complex multiscale biological processes spanning from tissue-level to cellular and molecular levels [1–3]. Recent advancements in microscopy and imaging technologies have provided unprecedented spatial and temporal resolution, enabling direct observation of complex biological phenomena under *in vitro, ex vivo*, and *in vivo* conditions [4–6]. These imaging modalities empower researchers to investigate dynamic cellular behaviors, protein trafficking, and cell–cell interactions, significantly deepening our understanding of fundamental biological mechanisms and disease pathophysiology [7, 8].

As the volume of biomedical images continues to grow, efficient and accurate image analysis has become increasingly important. Conventional methods for identifying, classifying, and tracking cells and proteins face several critical limitations. Classical approaches, such as intensity thresholding, edge detection, and region-based segmentation often struggle to accommodate the variability, deformation, and motion inherent in biological imaging, especially in live-cell experiments [9–11]. Manual annotation, although highly accurate when performed by trained experts, is labor-intensive, subjective, and impractical for large-scale or real-time applications [12, 13]. Furthermore, traditional image-processing pipelines typically rely on domain-specific parameters that require extensive manual tuning, and their performance degrades rapidly under conditions of low contrast, complex cellular morphology, or nonuniform illumination [14, 15].

To overcome these limitations, the biomedical research community has increasingly turned to machine learning and deep learning [16, 17]. Before the advent of deep learning, object detection relied on sliding window strategies and handcrafted features such as Histogram of Oriented Gradients (HOG) [18] and Scale-Invariant Feature Transform (SIFT) [19]. The introduction of convolutional neural networks (CNNs) subsequently enabled more capable detectors, including R-CNN, Fast R-CNN, and Faster R-CNN [20, 21], though these models remained computationally intensive and unsuitable for real-time use. These limitations motivated the You Only Look Once (YOLO) framework [22], which reformulated detection as a single regression task and enabled real-time end-to-end inference. Later generations progressively improved detection through anchor boxes [23], multi-scale feature extraction [24], richer backbone architectures [25, 26], and multi-task extensions, including segmentation and pose estimation [27, 28]. More recent versions such as YOLO11 and YOLO12 [29] incorporated temporal modules and cross-modal fusion, while YOLO26 [30] achieved NMS-free deployment with up to 43% reduction in CPU inference time. In parallel, Detection Transformers [31] brought attention-based architectures to object detection, with variants such as RT-DETR and RF-DETR offering competitive accuracy at reduced latency.

Despite these advances, applying joint detection-classification models to biomedical imaging tasks that require fine-grained morphological discrimination remains challenging. Biomedical images differ substantially from natural-scene images used for pretraining: they typically exhibit lower spatial resolution, high background noise, uneven illumination, and poor contrast between objects and surrounding media [4, 5, 32]. Beyond these image-quality issues, biomedical classification often requires distinguishing morphological variants within the same object category based on subtle differences in shape, surface texture, or intensity distribution [33–35]. Joint detection-classification architectures address both tasks through shared feature representations that are primarily shaped by bounding box regression objectives. Because effective localization requires features that are invariant to object position and scale, the learned representations tend to suppress the position-sensitive textural cues that fine-grained classification depends on. This tension is further aggravated by severe class imbalance: when minority phenotypes constitute only a small fraction of training instances, they contribute insufficient gradient signal to shape the shared backbone, and standard imbalance-mitigation strategies applied at the data or loss level may not adequately compensate for this architectural mismatch.

These challenges are particularly evident in sickle cell disease (SCD), where accurate phenotypic characterization of red blood cells (RBCs) is essential for understanding disease heterogeneity and evaluating therapeutic response [36–47]. SCD RBCs present across five morphologically distinct categories: discocytes (DO), echinocytes (E), elongated and sickle-shaped cells (ES), granular cells (G), and reticulocytes (R). Although AI-based classifiers have been developed for automated phenotyping [48–53], most are trained on pre-cropped, single-cell images. This approach bypasses the complexity of real microscopy images, where cells are densely packed and frequently overlapping, and where class imbalance is severe: the dominant DO class accounts for over 70% of all instances while rare phenotypes such as R represent fewer than 2%. Whether joint detection-classification models can reliably handle these conditions remain an open question that existing benchmarks, built on isolated single-cell crops, do not address.

In this work, we investigate this question systematically and propose a practical solution. We first benchmark a broad range of state-of-the-art joint detectors across multiple generations and scales of both YOLO and DETR on fully annotated full-scope microscopy images of SCD RBCs. We find that while these models achieve reliable cell localization, their classification performance for minority phenotypes falls well below that of specialized single-cell classifiers. We further evaluate five imbalance-mitigation strategies, including copy-paste augmentation, focal loss, class weighting, rare-class cropping, and oversampling, and show that all yield only marginal improvements within joint architectures. These results suggest that the performance limitation is rooted in the architectural design of joint models rather than in data quantity alone.

Based on these findings, we propose a two-step framework that fully separates cell localization from morphological classification. A YOLO26n detector first identifies and localizes individual RBCs in full-scope images; each detected region is then cropped into a standardized single-cell patch and passed to a fine-tuned DenseNet121 ensemble trained exclusively for morphological discrimination. Because the classifier operates on position-normalized patches without any localization objective, it can focus entirely on the textural and boundary features that distinguish minority phenotypes, rather than sharing capacity with the detection task. The main contributions of this work are as follows:

1. **Systematic benchmarking of joint detectors on full-scope SCD images**. We provide the first comprehensive evaluation of 11 model variants from the YOLO and DETR families on fully annotated full-scope microscopy images, demonstrating that all tested models show significant performance degradation for minority phenotypes (e.g., G: mAP@50 as low as 0.095; R: mAP@50 as low as 0.204), regardless of architecture or scale.
2. **Evaluation of imbalance-mitigation strategies within joint architectures**. We benchmark five commonly used imbalance-handling strategies on this task and show that none produces meaningful improvements for the most challenging minority classes, providing evidence that data-level augmentation alone is insufficient when the underlying architecture is not suited for fine-grained classification.
3. **A two-step decoupled framework for full-scope RBC phenotyping**. The proposed pipeline achieves a macro-average F1 improvement of +0.1675 over the single-step baseline, with particularly large gains for clinically important minority phenotypes (G: +0.2774; R: +0.2603; E: +0.1623).
4. **A curated full-scope evaluation dataset and protocol**. We establish an annotated evaluation protocol based on 497 full-scope microscopy images comprising 30,991 cells from 8 patients across 2 institutions, enabling systematic assessment of RBC detection and phenotype classification under realistic imaging conditions that pre-cropped single-cell datasets cannot capture.

## 2 Data and Methods

### 2.1 Dataset

We utilize a sickle RBC image dataset originally collected in Xu et al. [48], which comprises 497 high-resolution full-scope microscopy images of 8 different SCD patients collected from two different hospitals, as shown in the Figs. 1(A-B).

**Figure 1:**
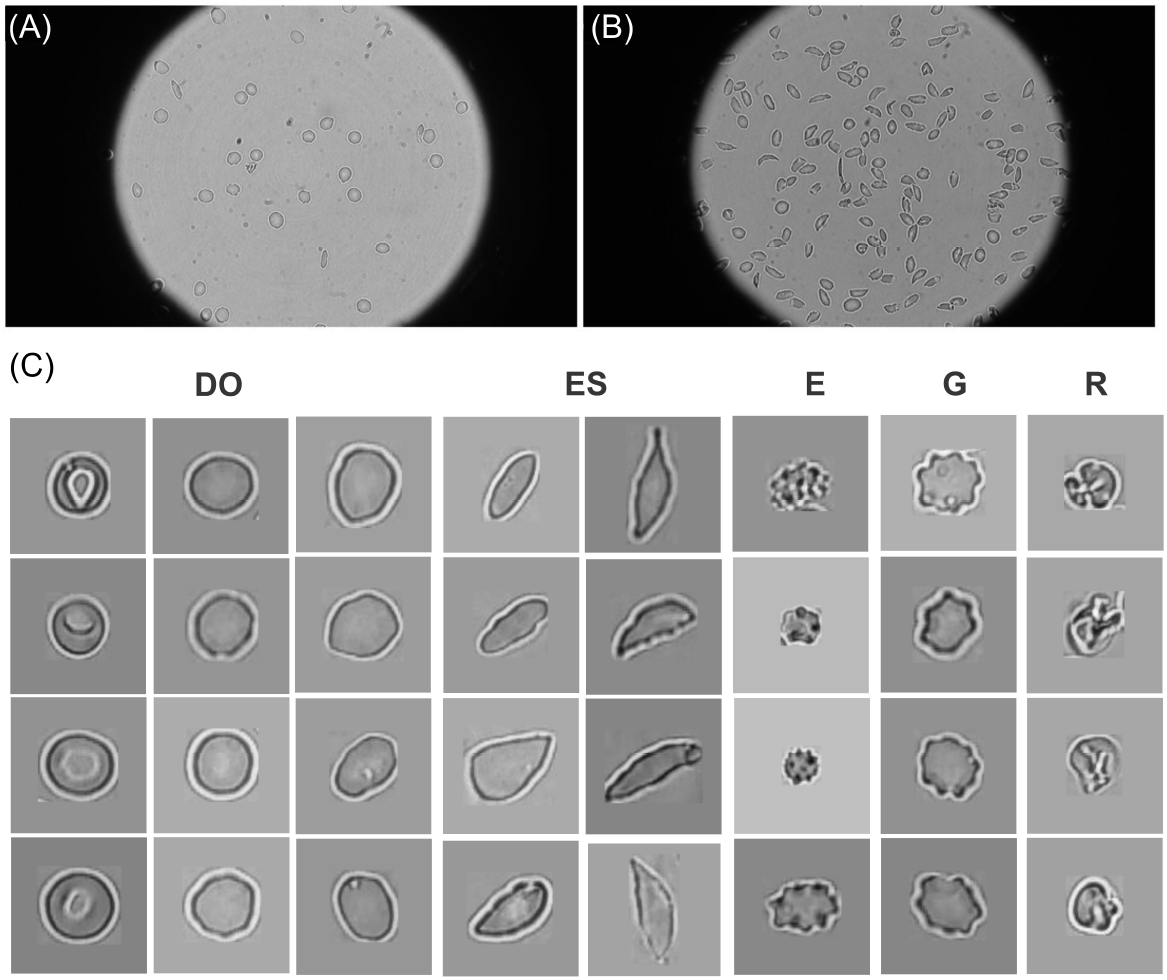
Images of sickle RBCs captured under full-scope bright-field microscopy from Xu et al. [48], showing a relatively sparse (A) and dense (B) population of RBCs. (C) Representative single-cell images of the five-class RBC phenotypes, cropped from full-scope microscopic images (A-B). The five phenotypes include DO (Discocytes & Ovalocytes), ES (Elongated & Sickle cells), E (Echinocytes), G (Granular cells), and R (Reticulocytes).

Each image was acquired using a Zeiss inverted microscope with a 63× oil immersion objective and a Sony Exmor CMOS sensor, at a resolution of 1920×1080 pixels. Blood samples were obtained from SCD patients through a standard RBC density fractionation protocol and imaged *ex vivo* under normoxic conditions. From these images, we extracted and segmented 31,297 individual RBC images, covering a wide spectrum of morphological variations present in the raw data. As illustrated in the Fig. 1(C), we categorized these RBCs into five morphological phenotypes, namely, DO (Discocytes & Ovalocytes), ES (Elongated & Sickle cells), E (Echinocytes), G (Granular cells), and R (Reticulocytes). Human cell-level annotation was conducted using the Roboflow platform [54] on 497 images, yielding a total of 30,991 annotated cells and facilitating corresponding YOLO-format label files for downstream detection and subtype classification. To ensure a fair evaluation, 100 images are used for model evaluation while the remaining 397 images are used to fine-tune detection models and the downstream subtype classifiers. As shown in Table 1, we crop 24,688 single-cell images from the 397 annotated images. As the number of RBCs in DO and ES classes greatly outnumber other subtypes, we apply random downsampling to DO and ES to prevent overfitting. All cropped images are then subjected to manual quality control to remove noisy, low-quality, or mislabeled samples. After this refinement, as summarized in Table. 1, a final set of 3,079 high-quality single-cell images is used to train the classification models. Compared with the single-cell dataset reported in Xu et al. [48], which contained 1,004 pre-cropped images, our annotation covers the complete full-scope images and yields a substantially larger and more morphologically diverse set of 3,079 quality-controlled single-cell crops drawn from a wider range of imaging conditions across the 8 patients. (see Table 1), resulting in more comprehensive morphological coverage. This diversity supports improved model generalization and robustness.

**Table 1:**
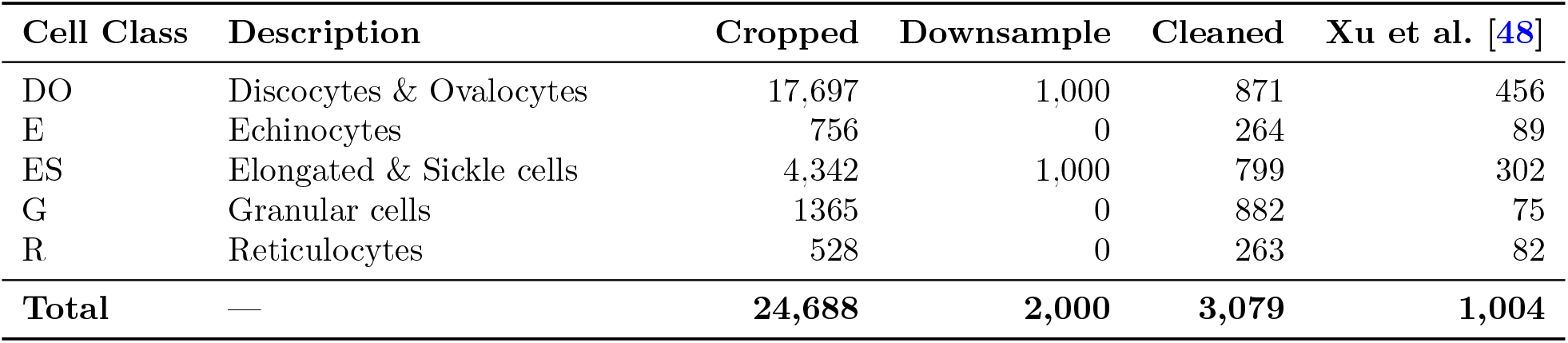
Number of manually labeled single-cell images across RBC classes cropped from the 397 full-scope images.

### 2.2 Detection and classification using YOLO and DETR

We first benchmark end-to-end detection–classification performance using two representative families of modern vision detectors, namely YOLO-based detectors and DETR-style transformer detectors. All microscopy images are annotated in COCO format with five morphological categories, where DO is the dominant class and four rarer classes include E, ES, G, and R. Specifically, for the YOLO family, we evaluate multiple generations and scales, including YOLO11, YOLO12, and YOLO26, as well as the newly released multi-scale YOLO26 variants. For the DETR family, we evaluate RT-DETR and RF-DETR models, including RT-DETR-R50 and RF-DETR in nano/small/medium sizes.

Despite strong object localization performance, fine-grained morphological classification remains challenging in microscopy due to subtle inter-class differences (e.g., between E and R) and severe class imbalance, with DO accounting for the vast majority of instances while R is scarce. As a reference, we train each detector on the original training set without any re-weighting, re-sampling, or targeted augmentation. In all subsequent experiments, we keep the model architectures and training hyperparameters fixed and vary only the imbalance-handling strategy, enabling fair evaluation of the performance of the proposed mitigation methods.

#### 2.2.1 Imbalance-mitigation strategies

To address the imbalance-induced degradation in rare-class recognition while maintaining performance on the dominant DO class, we evaluate complementary approaches at both the loss level and the data level. At the loss level, we apply class-weighted cross-entropy and focal loss [55]. At the data level, we apply image-level oversampling, targeted copy–paste augmentation [56], and rare-class cropping. These approaches are introduced as follows.

##### Class-Weighted Loss

We increase the learning signal from rare categories by applying class weights to the classification loss:

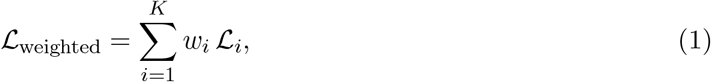

where *K* is the number of classes, ℒ_*i*_ denotes the loss contribution for class *i*, and *w*_*i*_ is computed from class frequency. We compare three commonly used weighting schemes:

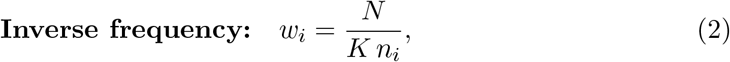

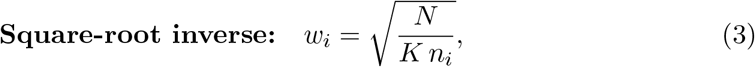

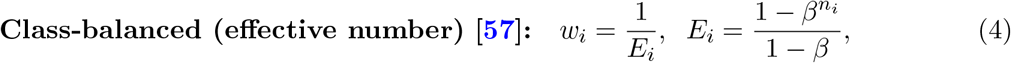

where *N* denotes the total number of samples, *n*_*i*_ is the number of samples for class *i*, and *β* is set to 0.9999.

##### Focal loss

To down-weight easy examples and focus training on hard and/or rare cases, we also evaluate focal loss:

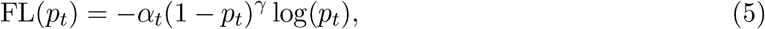

where *p*_*t*_ is the predicted probability of the ground-truth class, *α*_*t*_ is the class balancing factor, and *γ* is the focusing parameter that reduces the loss contribution from easy examples. We set *α*_*t*_ = 0.25 and *γ* = 2.0.

##### Image-level oversampling

We oversample images that contain at least one rare-class instance (E/ES/G/R) by duplicating such images in the training set, as illustrated in Figs. 2(A-B). Let *N*_maj_ denote the majority-class count and *N*_rare_ the total number of rare-class instances. Given a target ratio *r*, the duplication factor is

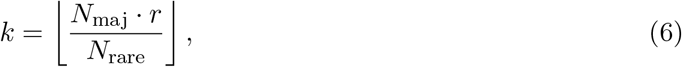

with image and annotation identifiers reassigned to preserve COCO consistency. Unless stated otherwise, we set *r* = 0.5.

**Figure 2:**
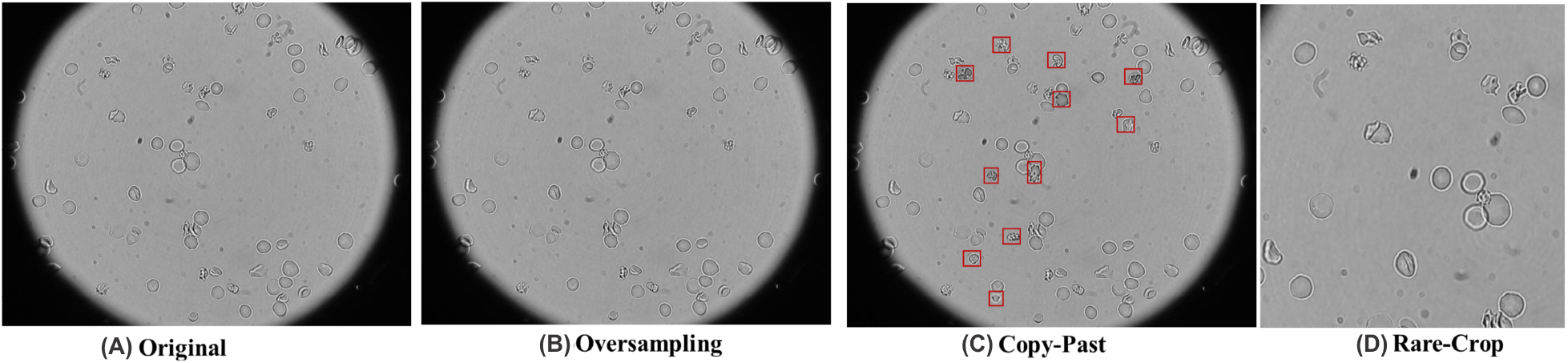
Three strategies to mitigate class imbalance of sickle RBCs captured in full-scope images. (A) Original microscopy scope exhibiting an imbalanced class distribution. (B) Image-level over-sampling increases the sampling frequency of images containing rare morphologies without altering pixel content. (C) Targeted copy–paste augmentation inserts rare-cell patches into feasible locations; red boxes indicate pasted instances. (D) Rare-class cropping produces centred crops from full-scope images around rare instances to increase their visual prominence during training.

##### Targeted copy–paste augmentation

To increase both the prevalence and visual diversity of rare morphologies, we implement targeted copy–paste augmentation. As illustrated in Fig. 2(C), rare-class patches are extracted from annotated bounding boxes with a 5-pixel margin and pasted into feasible locations within the field of view. Candidate paste locations are constrained to fall within an approximate circular imaging region (radius 0.45 times the image size) and to avoid excessive overlap with existing objects (IoU < 0.3). Pasted patches are blended using an elliptical soft mask *M* smoothed by a Gaussian filter:

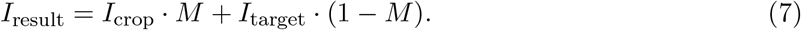

Unless stated otherwise, copy–paste is applied to all training images (augment ratio = 1.0) while retaining the original images.

##### Rare-class cropping

To increase the prominence of rare instances and encourage fine-grained feature learning, we generate additional crops centred on each rare-class instance, as shown in Fig. 2(D). For a rare instance with centre (*c*_*x*_, *c*_*y*_), we produce *k* = 2 crops of size 640 × 640 with random centre offsets

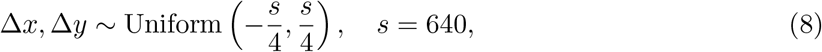

and transform all annotations within each crop accordingly. Original full-frame images are retained to preserve contextual diversity.

### 2.3 Two-step pipeline for sickle RBC detection and classification

Although YOLO and DETR models can detect and classify objects concurrently, the performance summarized in Table 2 indicates that, even with proper fine-tuning, these models reliably detect RBCs with diverse morphologies in microscopic images, whereas their performance in fine-grained sickled-RBC subtype classification remains suboptimal. This is likely because such models adopt a detection-oriented design optimized for multi-object localization and real-time inference, which can be less effective at capturing subtle intra-class morphological differences required for fine-grained phenotype recognition. Therefore, we propose a two-step pipeline that combines YOLO or DETR’s strengths in detection with specialized image classifiers for detailed RBC morphology classification. The overall workflow is illustrated in Fig. 3.

**Table 2:** Per-class one-step detection and classification performance across various YOLO and DETR models. The best results within each class block are highlighted in bold.

| Class | Images | Instances | Model | Precision | Recall | mAP@50 | mAP@50:95 |
| --- | --- | --- | --- | --- | --- | --- | --- |
| DO | 80 | 4354 | YOLO11 | 0.908 | **0.974** | 0.984 | 0.867 |
|  |  |  | YOLO12 | 0.888 | 0.959 | 0.976 | 0.829 |
|  |  |  | RT-DETR (R50) | 0.879 | 0.934 | 0.947 | 0.773 |
|  |  |  | RF-DETR Nano | 0.941 | 0.928 | 0.993 | 0.905 |
|  |  |  | RF-DETR Small | 0.955 | 0.959 | **0.996** | **0.932** |
|  |  |  | RF-DETR Medium | **0.960** | 0.957 | 0.964 | 0.8599 |
|  |  |  | YOLO26n | 0.916 | 0.946 | 0.979 | 0.875 |
|  |  |  | YOLO26s | 0.923 | 0.967 | 0.986 | 0.896 |
|  |  |  | YOLO26m | 0.943 | 0.964 | 0.987 | 0.904 |
|  |  |  | YOLO26l | 0.916 | 0.946 | 0.979 | 0.875 |
|  |  |  | YOLO26x | 0.954 | 0.952 | 0.987 | 0.910 |
| E | 56 | 207 | YOLO11 | 0.723 | 0.773 | 0.813 | 0.593 |
|  |  |  | YOLO12 | 0.502 | 0.585 | 0.540 | 0.382 |
|  |  |  | RT-DETR (R50) | 0.598 | 0.425 | 0.417 | 0.276 |
|  |  |  | RF-DETR Nano | 0.784 | 0.527 | 0.879 | 0.696 |
|  |  |  | RF-DETR Small | 0.852 | 0.778 | **0.906** | **0.771** |
|  |  |  | RF-DETR Medium | 0.858 | 0.729 | 0.887 | 0.680 |
|  |  |  | YOLO26n | 0.719 | 0.725 | 0.791 | 0.588 |
|  |  |  | YOLO26s | 0.774 | **0.826** | 0.873 | 0.686 |
|  |  |  | YOLO26m | 0.833 | 0.773 | 0.902 | 0.737 |
|  |  |  | YOLO26l | 0.719 | 0.725 | 0.791 | 0.588 |
|  |  |  | YOLO26x | **0.872** | 0.786 | 0.904 | 0.731 |
| ES | 76 | 796 | YOLO11 | 0.737 | 0.803 | 0.850 | 0.705 |
|  |  |  | YOLO12 | 0.488 | **0.843** | 0.737 | 0.576 |
|  |  |  | RT-DETR (R50) | 0.712 | 0.704 | 0.682 | 0.495 |
|  |  |  | RF-DETR Nano | 0.815 | 0.680 | 0.940 | 0.810 |
|  |  |  | RF-DETR Small | **0.843** | 0.769 | **0.959** | **0.865** |
|  |  |  | RF-DETR Medium | 0.833 | 0.784 | 0.889 | 0.753 |
|  |  |  | YOLO26n | 0.726 | 0.788 | 0.821 | 0.689 |
|  |  |  | YOLO26s | 0.738 | 0.822 | 0.863 | 0.748 |
|  |  |  | YOLO26m | 0.768 | 0.840 | 0.890 | 0.781 |
|  |  |  | YOLO26l | 0.726 | 0.788 | 0.821 | 0.689 |
|  |  |  | YOLO26x | 0.805 | 0.806 | 0.893 | 0.783 |
| G | 47 | 139 | YOLO11 | 0.436 | 0.517 | 0.522 | 0.430 |
|  |  |  | YOLO12 | 0.139 | 0.014 | 0.095 | 0.076 |
|  |  |  | RT-DETR (R50) | 0.374 | 0.216 | 0.187 | 0.156 |
|  |  |  | RF-DETR Nano | **0.849** | 0.324 | **0.933** | 0.815 |
|  |  |  | RF-DETR Small | 0.705 | 0.482 | 0.882 | **0.825** |
|  |  |  | RF-DETR Medium | 0.717 | 0.547 | 0.660 | 0.576 |
|  |  |  | YOLO26n | 0.438 | 0.594 | 0.497 | 0.415 |
|  |  |  | YOLO26s | 0.659 | 0.488 | 0.607 | 0.536 |
|  |  |  | YOLO26m | 0.608 | **0.676** | 0.685 | 0.603 |
|  |  |  | YOLO26l | 0.438 | 0.594 | 0.497 | 0.415 |
|  |  |  | YOLO26x | 0.589 | 0.612 | 0.701 | 0.609 |
| R | 43 | 105 | YOLO11 | 0.533 | 0.467 | 0.452 | 0.335 |
|  |  |  | YOLO12 | 0.234 | 0.399 | 0.204 | 0.132 |
|  |  |  | RT-DETR (R50) | 0.317 | 0.286 | 0.243 | 0.161 |
|  |  |  | RF-DETR Nano | 0.595 | 0.238 | 0.655 | 0.593 |
|  |  |  | RF-DETR Small | 0.638 | 0.486 | **0.738** | **0.650** |
|  |  |  | RF-DETR Medium | 0.654 | 0.505 | 0.626 | 0.487 |
|  |  |  | YOLO26n | 0.480 | 0.543 | 0.509 | 0.386 |
|  |  |  | YOLO26s | 0.527 | 0.498 | 0.511 | 0.402 |
|  |  |  | YOLO26m | 0.513 | **0.590** | 0.567 | 0.451 |
|  |  |  | YOLO26l | 0.480 | 0.543 | 0.509 | 0.386 |
|  |  |  | YOLO26x | **0.703** | 0.562 | 0.695 | 0.560 |

**Figure 3:**
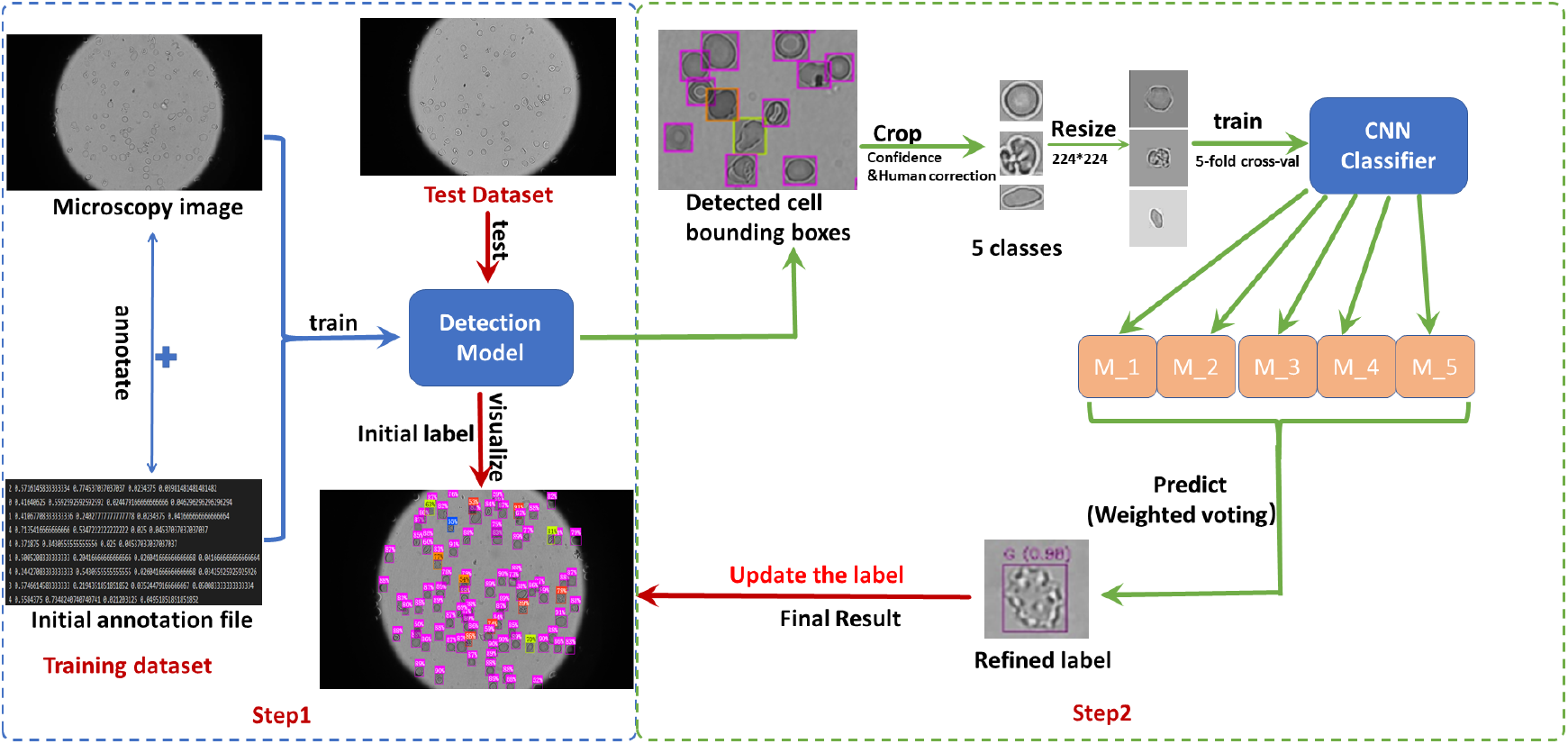
Overview of the proposed two-step pipeline for the detection and classification of sickle RBCs within full-scope microscopic images. The detection model first localizes and crops individual cells from full-scope microscopy images, followed by a fine-tuned classifier that assigns each cell to one of five morphological categories. The labels will be passed back to the full-scope images for visualization.

#### Step 1: RBC detection

Full-scope microscopy images in the training dataset are manually annotated with bounding boxes corresponding to individual RBCs. These annotations are used to fine-tune the YOLO or DETR-based model [29] for efficient detection of RBCs with various shapes. The trained model is then applied to the test dataset to localize RBCs. Detected bounding boxes with confidence scores are used to crop cell regions, which are subsequently resized to a standardized input resolution for downstream classification.

#### Step 2: Morphological phenotype classification using AI-based classifiers

In the second step, classifiers trained on single-cell images are used to categorize each detected RBC into one of five predefined morphological classes (Fig. 1C). We benchmark five modern architectures: ResNet18 [58], ResNet50 [58], DenseNet12 [59], EfficientNet-B3 [60], and Vision Transformer (ViT) [61]. To improve generalization and robustness to pose and spatial variation while preserving morphology-critical features, a lightweight geometric augmentation pipeline is applied exclusively to the training data. Each training image undergoes random horizontal and vertical flipping, small-angle rotation (constrained to *±*12^◦^), and mild affine perturbation comprising limited translation (up to 5% of image size) and isotropic scaling (within [0.95, 1.05]). No photometric augmentation is applied, ensuring that the classifier focuses on stable morphological cues rather than appearance artifacts. Validation data remains unaugmented to ensure unbiased model selection. Class imbalance is addressed using weighted cross-entropy with inverse-frequency class weights computed from the training distribution within each fold. All classifiers are initialized from ImageNet pre-trained weights and fine-tuned end-to-end using AdamW (learning rate 1 × 10^−4^, weight decay 1 × 10^−4^). The learning rate is adjusted via ReduceLROnPlateau when validation loss plateaus (patience = 3, factor = 0.5, minimum learning rate = 1 × 10^−6^). Early stopping is applied based on validation performance (patience = 7, *δ* = 5 × 10^−4^), retaining the best checkpoint for inference. To improve robustness under limited training data, all classifiers are trained and evaluated using 5-fold cross-validation. For each RBC, the final morphological label is determined by confidence-weighted ensemble voting over the five fold-specific models, where each model contributes its maximum soft-max confidence to the predicted class score and the class with the largest accumulated score is assigned as the final prediction.

### 2.4 Evaluation metrics

We evaluate the proposed framework from three aspects: (i) cell-level morphology classification, (ii) object detection, and (iii) computational efficiency. The results are reported as the mean across splits.

#### Classification metrics

Classification performance is evaluated using accuracy and averaged precision, recall, and F1 score under five-fold cross-validation. For each class *c*, we adopt a one-vs-rest formulation to compute *TP*_*c*_, *FP*_*c*_, *FN*_*c*_, as follows,

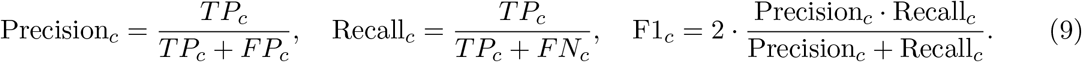

Accuracy is computed as the fraction of correctly classified samples across all classes,

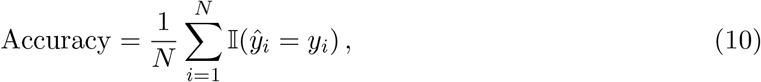

where *y*_*i*_ and *ŷ*_*i*_ denote the ground-truth and predicted labels for sample *i, N* is the total number of samples, and 𝕀 (·) is the indicator function. 𝕀 (·) returns 1 if its argument is true and 0 otherwise.

Macro-averaged metrics are obtained by averaging over the *K*=5 classes, for example

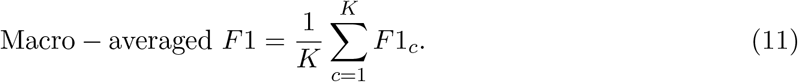

In imbalanced settings, *F*1_macro_ complements accuracy by weighting all classes equally and therefore better reflects balanced performance across both frequent and rare categories.

#### Detection metrics

Detection performance is reported using COCO-style average precision (AP) and mean AP (mAP). For each class, AP is computed from the precision–recall curve induced by sweeping the confidence threshold. We report mAP@0.5 (mAP@50) and mAP@0.5:0.95, which averages AP over IoU thresholds from 0.50 to 0.95 in steps of 0.05, following the standard COCO protocol.

#### Computational efficiency

We report inference throughput (frames per second; FPS) and latency measured at batch size 1 with 640 × 640 inputs on an NVIDIA RTX 3090 GPU. Model complexity is quantified by the number of parameters and FLOPs.

## 3 Results

### 3.1 Performance of YOLO and DETR models on detecting the five phenotypes of sickle RBCs from full-scope images

#### 3.1.1 Impact of model architecture and size on its performance

First, we evaluate multiple versions and model sizes from both the YOLO and DETR families to provide a comprehensive performance comparison. Specifically, we test two standard YOLO versions (YOLO11 and YOLO12), one real-time transformer-based model (RT-DETR with a ResNet-50 backbone), and three scaled variants of RF-DETR (Nano, Small, and Medium). In addition, we assess five different capacity levels of the YOLO26 architecture, including YOLO26n (Nano), YOLO26s (Small),YOLO26m (Medium), YOLO26l (Large), YOLO26x (Extra-Large). By evaluating both different model families and multiple scaling variants within the same architecture, we systematically analyze the effects of architectural design (CNN-based versus transformer-based detectors) and model capacity (from lightweight to high-capacity configurations) on detecting the five sickle RBC phenotypes.

Table 2 presents the per-class one-step detection and classification performance of various YOLO- and DETR-based models across five classes (DO, E, ES, G, and R), evaluated using Precision, Recall, mAP@50, and mAP@50:95. The dataset distribution varies notably among classes, with DO having the largest number of instances (4354 across 80 images), followed by ES (796 instances), E (207), G (139), and R (105), indicating a clear class imbalance. For the DO class, all models achieve strong performance, reflecting the large number of training instances. RF-DETR Small attains the best overall detection accuracy with the highest mAP@50 (0.996) and mAP@50:95 (0.932), while RF-DETR Medium achieves the highest precision (0.960). YOLO11 provides the highest recall (0.974). The YOLO26 variants also perform competitively, particularly YOLO26x, which achieves 0.954 precision and 0.910 mAP@50:95. In the E class, performance is comparatively lower, likely due to the less appearance compared to the RBCs within DO type. RF-DETR Small again achieves the best overall localization accuracy (mAP@50 = 0.906, mAP@50:95 = 0.771), while YOLO26x records the highest precision (0.872) and YOLO26s achieves the highest recall (0.826). YOLO12 and RT-DETR (R50) show notably weaker results in this class. For the ES class, RF-DETR Small consistently outperforms other models, achieving the highest precision (0.843), mAP@50 (0.959), and mAP@50:95 (0.865). YOLO12 achieves the highest recall (0.843) but with relatively low precision, indicating a tendency toward over-detection. The YOLO26 family, especially YOLO26m and YOLO26x, demonstrates balanced and competitive performance. The G class presents more challenging detection results due to the limited number of instances (139). RF-DETR Nano achieves the highest precision (0.849) and mAP@50 (0.933), while RF-DETR Small attains the best mAP@50:95 (0.825). YOLO26m achieves the highest recall (0.676), suggesting improved sensitivity but comparatively lower localization precision. YOLO12 performs poorly in this category. Similarly, the R class, with only 105 instances, shows modest performance across models. RF-DETR Small achieves the best overall detection accuracy (mAP@50 = 0.738, mAP@50:95 = 0.650), while YOLO26x achieves the highest precision (0.703). YOLO26m records the highest recall (0.590), indicating better object coverage at the cost of precision.

Notably, there is a significant drop in both precision and recall for the minority phenotypes (E, G, and R) compared to the majority class (DO). This decline suggests that model performance deteriorates as the number of training instances decreases, highlighting the impact of class imbalance on accurate detection and classification of less-represented phenotypes using YOLO and DETR models.

#### 3.1.2 Improving minor-class detection with imbalance-mitigation strategies

Since the class imbalance is the primary factor limiting morphology detection in full-scope image analysis, in this section, we investigate several imbalance mitigation strategies, including, copy–paste augmentation, focal loss, class weighting, rare-crop augmentation, and oversampling, to evaluate whether they can improve detection performance. Table 3 presents the per-class detection performance of different imbalance mitigation strategies applied to the RF-DETR Medium model. Altogether, the results indicate that while the majority class (DO) remains relatively stable across all methods, the minority classes (E, G, and R) exhibit more noticeable performance variations depending on the strategy used.

**Table 3:** Impact of imbalance mitigation approaches on per-class detection performance tested on the RF-DETR medium model. Metrics are reported in percent. Best performance in each column is shown in bold.

| Method | DO |  | E |  | ES |  | G |  | R |  |
| --- | --- | --- | --- | --- | --- | --- | --- | --- | --- | --- |
|  | mAP@50 | mAP@50:95 | mAP@50 | mAP@50:95 | mAP@50 | mAP@50:95 | mAP@50 | mAP@50:95 | mAP@50 | mAP@50:95 |
| copy_paste | 96.35 | **86.24** | 88.87 | 67.08 | 87.24 | 74.01 | 66.14 | 56.70 | **66.76** | **51.13** |
| focal_loss | 96.40 | 86.10 | 88.08 | 67.58 | 88.36 | 74.86 | **66.76** | **58.05** | 63.38 | 49.58 |
| baseline | **96.42** | 85.99 | 88.75 | 67.98 | **88.98** | **75.30** | 66.04 | 57.65 | 62.63 | 48.70 |
| class_weight | 96.39 | 85.96 | **88.96** | **68.07** | 88.34 | 74.84 | 66.14 | 57.45 | 61.77 | 47.97 |
| rare_crop | 96.14 | 85.33 | 85.84 | 64.96 | 85.57 | 71.91 | 61.97 | 53.21 | 56.69 | 44.42 |
| oversampling | 95.91 | 86.08 | 83.04 | 64.89 | 83.96 | 71.58 | 62.10 | 54.45 | 56.78 | 44.92 |

For the DO class, all approaches achieve very similar results, with mAP@50 values around 96%. The baseline model slightly outperforms others at mAP@50 (96.42%), while the copy-paste method achieves the highest mAP@50:95 (86.24%). This consistency suggests that imbalance mitigation techniques have limited impact on the well-represented majority class.

In the E class, modest improvements are observed with class-weight, which achieves the best performance at both mAP@50 (88.96%) and mAP@50:95 (68.07%). The baseline and focal-loss approaches also perform competitively, while rare-crop and oversampling result in noticeable declines. This indicates that moderate reweighting strategies are more effective than aggressive resampling for this minority phenotype for our dataset.

For the ES class, the baseline model delivers the best performance (88.98% mAP@50 and 75.30% mAP@50:95), suggesting that additional imbalance handling does not significantly improve results for this moderately represented class. Other methods produce comparable but slightly lower scores.

Greater differences appear in the more challenging G and R classes. In the G class, focal-loss achieves the highest performance (66.76% mAP@50 and 58.05% mAP@50:95), slightly outperforming other strategies. For the R class, copy-paste provides the most substantial improvement, reaching the highest mAP@50 (66.76%) and mAP@50:95 (51.13%), compared to the baseline (62.63% and 48.70%). In contrast, rare-crop and oversampling consistently underperform across minority classes, indicating that simple resampling or cropping strategies may introduce noise or reduce generalization.

To further evaluate the impact of minority-class performance on overall detection effectiveness, we present a minority trade-off plot in Fig. 4, which visualizes the relationship between overall mAP@0.5 and the average mAP@0.5 of the minority classes. This analysis highlights how different imbalance mitigation strategies affect the balance between global performance and minority-class accuracy. The results show that copy–paste augmentation shifts the performance point toward the desirable top-right region of the plot, indicating improved minority performance without sacrificing overall detection accuracy. In contrast, oversampling and rare-crop move the performance point toward the lower-left region, reflecting degradation in both overall and minority performance. These findings further support the effectiveness of copy–paste as a balanced strategy for handling class imbalance.

**Figure 4:**
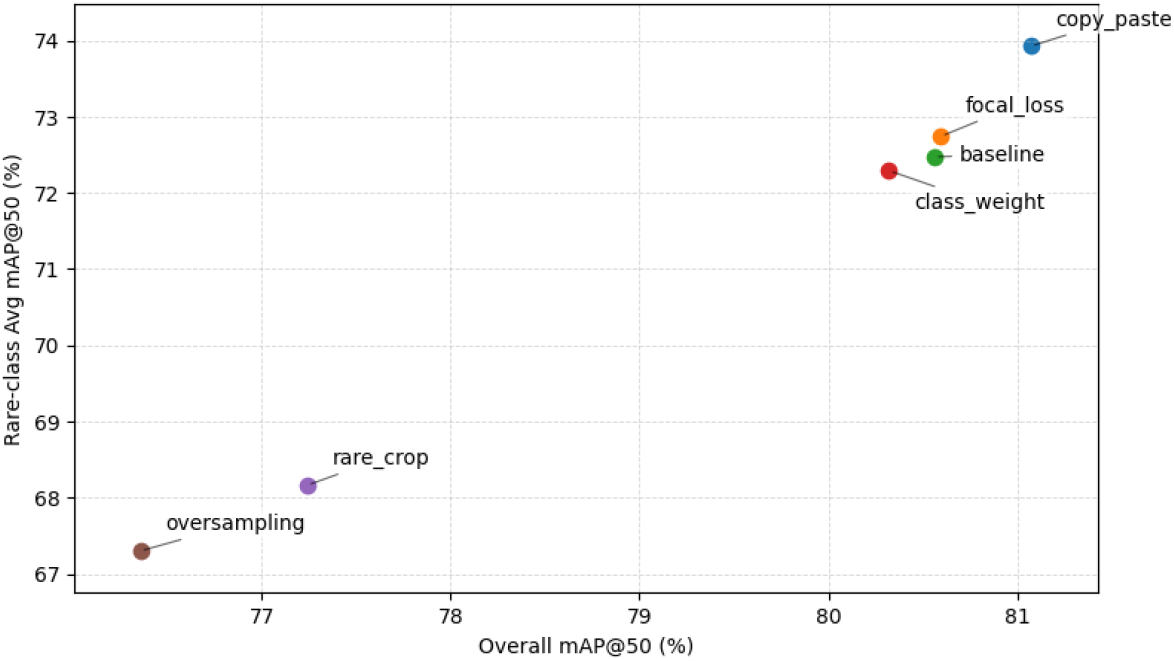
Trade-off between overall detection performance and minority-class accuracy across the five imbalance-mitigation strategies applied to RF-DETR Medium. Each point represents one strategy. The x-axis reports the overall mAP@0.5 across all five classes; the y-axis reports the average mAP@0.5 over the three minority phenotypes (E, G, and R). Strategies in the upper-right region achieves high performance on both overall and minority classes simultaneously. Copy-paste augmentation achieves the best trade-off, while oversampling and rare-crop degrade performance on both axes.

In summary, the findings indicate that although lightweight imbalance-aware techniques, such as focal loss and copy–paste augmentation, can improve performance for minority phenotypes without degrading performance on the majority class, the magnitude of these gains remains limited. This limited effect is likely attributable to the detection-oriented nature of these models. Architectures such as YOLO and DETR are primarily optimized for multi-object localization and real-time inference, emphasizing bounding box regression and object-level discrimination. As a result, they may be less effective at capturing the subtle intra-class morphological differences required for RBC phenotype recognition, particularly for rare categories with limited training samples.

### 3.2 Results of two-step identification and classification framework

The underperformance observed with YOLO and DETR models in classifying different RBC phenotypes motivates our proposed two-step framework, where RBCs are first localized within full-scope images and standardized through cropping, after which they are classified at the single-cell level, as illustrated in Fig. 3. The obtained labels will be sent back to update the label on the full-scope images. By decoupling localization from morphology discrimination, the framework allows the classification network to focus exclusively on detailed morphological features rather than jointly optimizing for detection and categorization, thereby improving the model’s ability to capture subtle differences within the five phenotypes of sickle RBCs.

To determine the detection model, we compare the class-agnostic localization accuracy and inference efficiency of three detectors. As shown in Table 4, YOLO26n achieves the highest Recall@0.75 (0.9830) with relatively low latency (8.26 ms/image), closely followed by YOLO11n (0.9795, 7.53 ms/image), whereas RF-DETR Nano attains a lower recall and substantially higher end-to-end latency (0.9199, 20.25 ms/image). The longer post-processing time of YOLO11n is mainly attributable to NMS, which is required to suppress redundant dense predictions; in contrast, YOLO26n and RF-DETR Nano employ an end-to-end, NMS-free design, resulting in faster post-processing. Moreover, RF-DETR Nano is transformer-based, and the attention/decoder computations incur higher inference overhead, leading to slower GPU inference than the convolution-dominated YOLO models. Altogether, considering both localization accuracy and runtime, YOLO26n is selected as the detector in our two-step detection-then-classification pipeline.

**Table 4:**
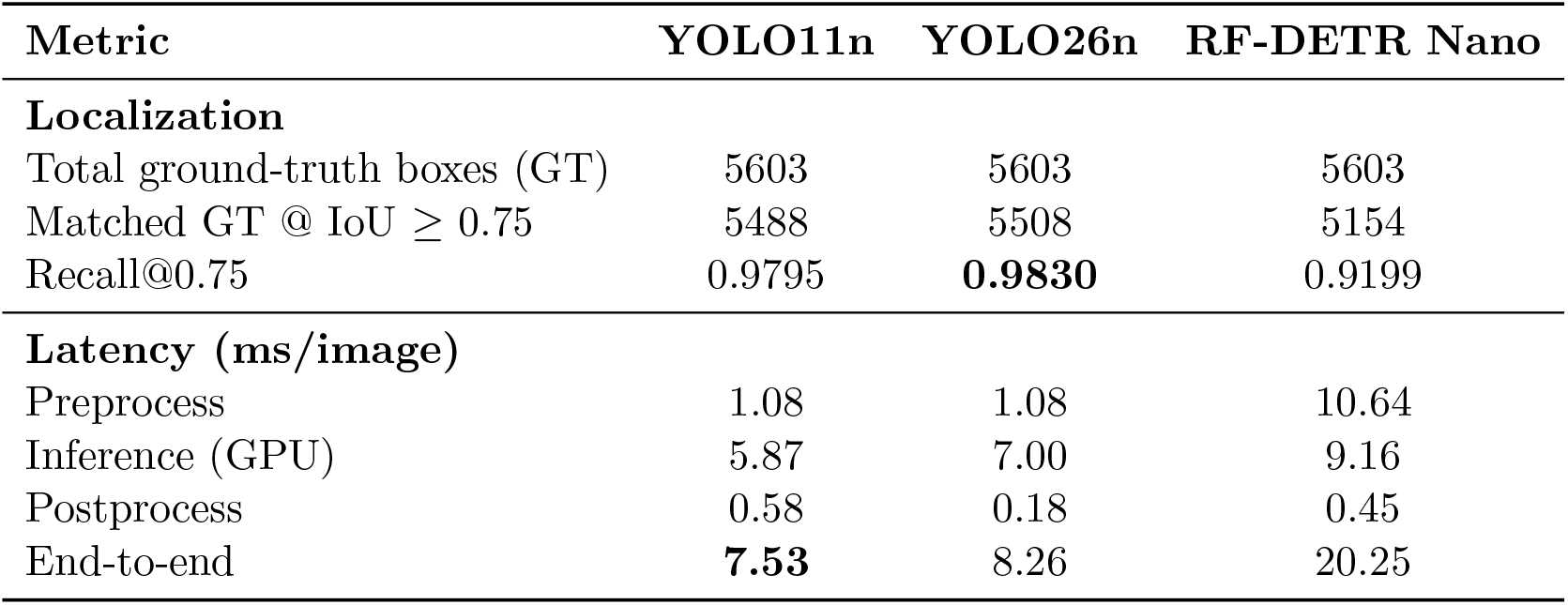
Performance and runtime efficiency of YOLO11n, YOLO26n and RF-DETR nano on detecting RBCs in full-scope images. Best results are highlighted in **bold**. Class-agnostic evaluation ignores phenotype labels and only assesses box matching using IoU. All timings are reported with input size 640 × 640 under identical hardware/software settings.

#### 3.2.1 Selection of the deep-learning based classifiers

In the second step, the cropped cell images from full-scope images using YOLO26n are fed into a pretrained classifier for subtype identification into: DO, E, ES, G, and R. To determine the most suitable architecture for our classifier, we test the performance of five deep learning models, namely Resnet18[62], ResNet50[58], DenseNet121[59], EfficientNet-B3[60], Vision Transformer (ViT)[61], on a five-class sickle RBC classification task. The performance of these models is summarized in Fig. 5 based on three evaluation metrics, including accuracy, recall and F1-score, which are averaged over five-fold cross-validation to ensure statistical reliability and robustness. These results show that while all models achieve strong performance, DenseNet121 outperforms the others across the three metrics.

**Figure 5:**
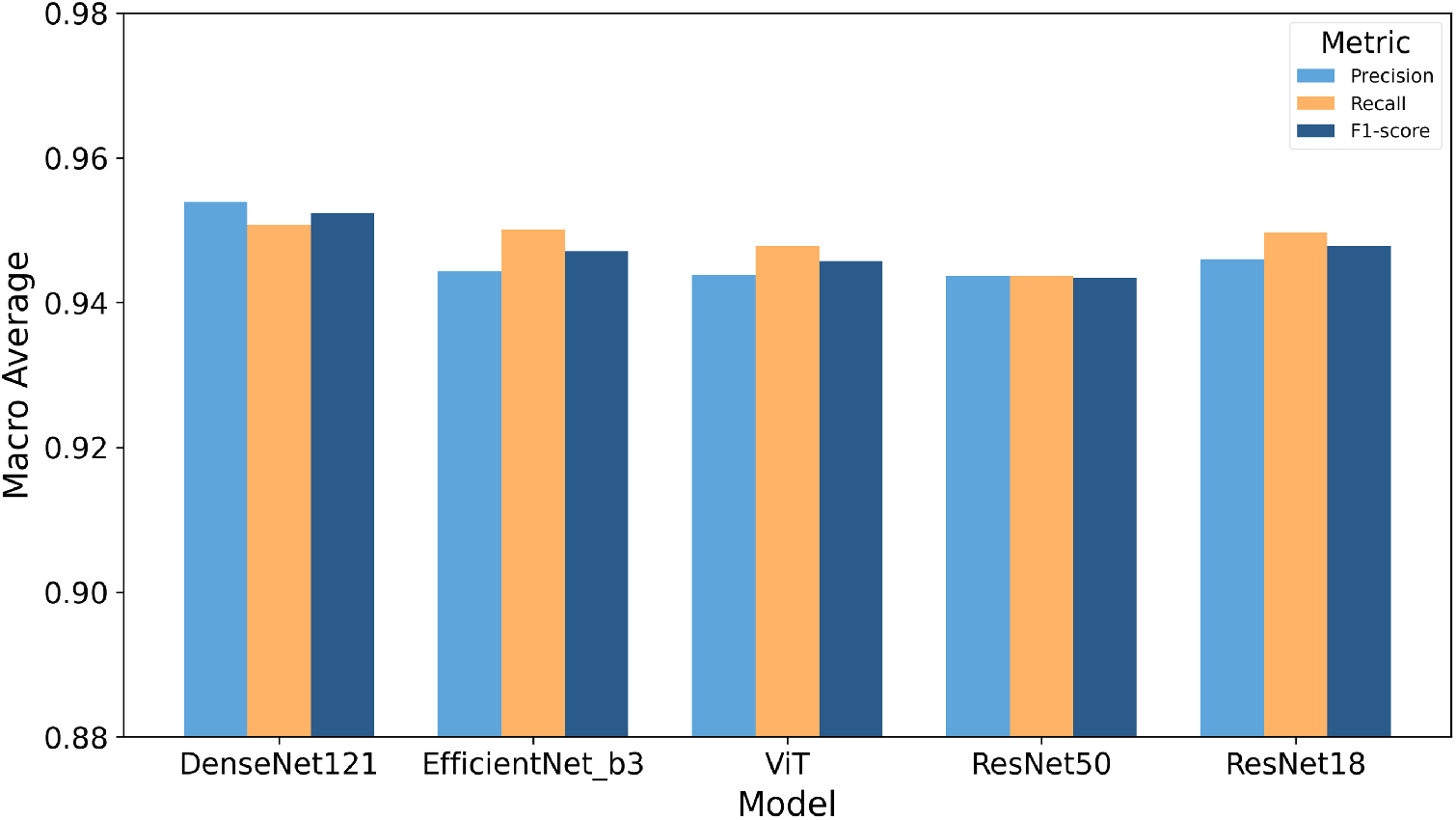
Performance comparison of five CNN-based and transformer-based models (DenseNet121, EfficientNet-B3, ViT, ResNet50, and ResNet18) in terms of macro-averaged precision, recall, and F1-score on the RBC morphology classification task, averaged over 5-fold cross-validation. DenseNet121 achieves the highest performance across all three metrics.

Table 5 reports the per-class F1-scores of all five models across the five RBC morphological categories. DenseNet121 achieves the highest F1-score in four out of five classes (DO, E, ES, and G), demonstrating its overall superiority for this task. ResNet18 obtains the best performance on class R (0.9336), marginally outperforming DenseNet121 (0.9333). Across all models, class E (Echinocytes) consistently yields the lowest F1-scores, ranging from 0.9151 (ResNet50) to 0.9272 (DenseNet121), suggesting that Echinocytes present greater morphological ambiguity and are more challenging to distinguish from other cell types. In contrast, classes ES and G achieve the highest F1-scores across all models, indicating more discriminative morphological features for these categories. ResNet50 shows the weakest overall performance, particularly on classes E (0.9151) and R (0.9162), while ViT achieves competitive results despite being the only non-CNN architecture evaluated, though it does not surpass the best CNN-based models in any class.

**Table 5:**
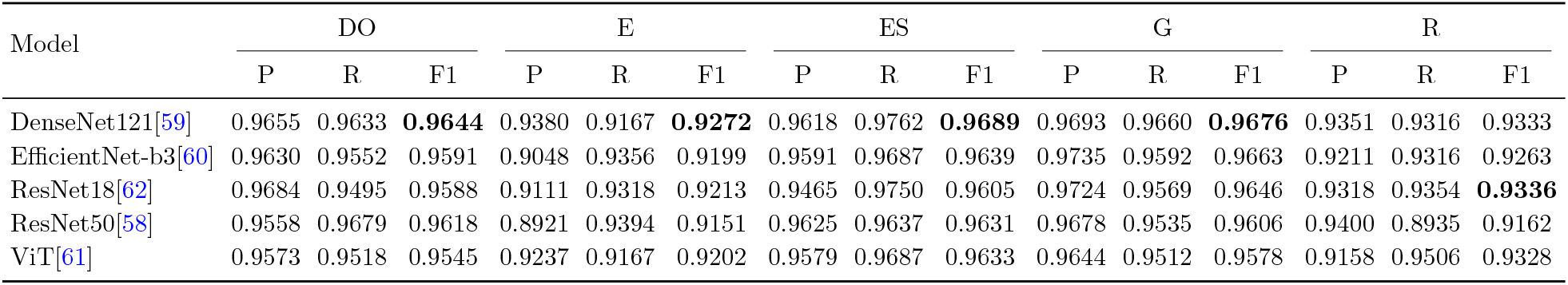
Performance comparison of five cell classification models (5-fold cross-validation average). Best values are bolded per-class F1. P/R/F1 denote precision/recall/F1-score.

#### 3.2.2 Analysis of misclassified cells by the classifier with the best performance

In this section, we further analyze the misclassified cells by the classifier with the best performance, DenseNet121, through which we can evaluate the challenges faced by the classification models. The confusion matrix in Fig. 6(A) shows strong diagonal dominance, indicating that most samples are correctly classified across all five classes (DO, E, ES, G, and R). The highest number of correct predictions is observed for DO (839) and ES (780), followed by G (852), E (242), and R (245). Misclassifications are relatively limited but reveal certain patterns. For example, DO samples are occasionally confused with ES (21) and G (8). This suggests that the classifier may be highly sensitive to elongation features, potentially overemphasizing shape deformation when distinguishing between normal biconcave morphology and elongated forms. Since both classes share elongated or oval characteristics under certain imaging conditions, subtle boundary distortions or orientation effects may contribute to this confusion. Class E shows some confusion with G (13) and R (9), suggesting feature similarity between these categories. Similarly, G instances are sometimes mis-classified as DO (14) and E (12). These results further support the hypothesis that surface texture irregularities are challenging for the classifier to differentiate consistently. Class R demonstrates minor confusion with ES (8) and E (4).

**Figure 6:**
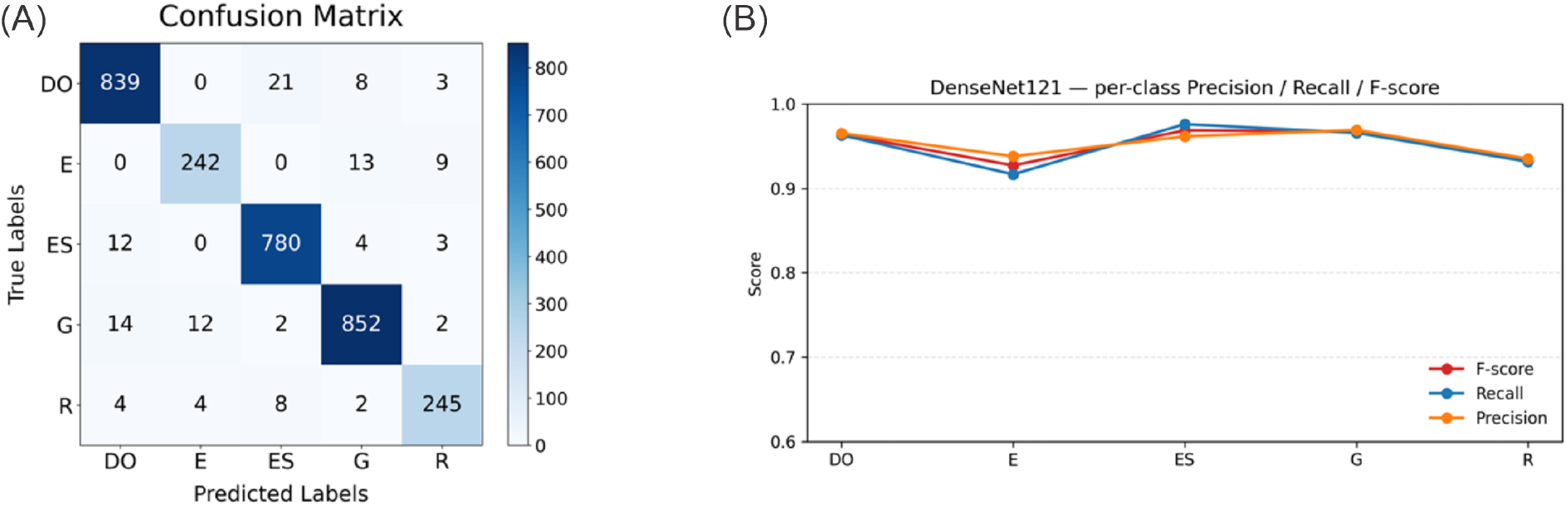
Performance of DenseNet121 on the five-class classification task. (A)Confusion matrix aggregated over 5-fold cross-validation, with rows denoting true labels and columns denoting predicted labels. (B) Per-class mean precision, recall, and F1-score across 5-fold cross-validation for classes DO, E, ES, G, and R.

Fig. 6(B) further quantifies the per-class performance of DenseNet121 using precision, recall, and F1-score. Classes DO and ES achieve the highest and most balanced scores, with F1 values of 0.9644 and 0.9689, respectively, confirming their strong separability observed in the confusion matrix. Class E yields the lowest scores across all three metrics (P = 0.9380, R = 0.9167, F1 = 0.9272), with recall notably lower than precision, indicating that a proportion of true E samples are misclassified into other categories, consistent with the confusion observed in Fig. 6(A). Class G maintains strong performance (F1 = 0.9676) with well-balanced precision and recall. Class R achieves an F1 of 0.9333, with precision and recall remaining comparable (P = 0.9351, R = 0.9316), suggesting no systematic bias toward false positives or false negatives. Overall, DenseNet121 demonstrates consistently high classification performance across all five classes, with the relatively lower performance on class E reflecting its greater inter-class similarity and higher misclassification rates identified in the confusion matrix.

#### 3.2.3 Performance evaluation of the two-step framework

In this section, we evaluate the performance of the proposed two-step framework on 100 full-scope microscopic images in the test dataset. The evaluation is conducted from both detection and classification perspectives. The detection results are summarized in Table 6. “Pred” denotes the total number of objects detected from the 100 test images. “TP” represents the number of correctly detected RBCs, “FP” indicates the number of objects incorrectly detected as RBCs, and “FN” denotes the number of missed RBCs. Based on these values, we compute Precision, Recall, and F1-score. As shown in Table 6, the proposed pipeline achieves detection-level performance with a Precision of 0.9873, Recall of 0.9458, and F1-score of 0.9661. These results demonstrate the detection model’s consistent performance in localizing individual RBCs across full-scope microscopic images. This also suggests that the two-step pipeline effectively leverages the advanced capabilities of YOLO models to detect objects in full-scope microscopic images containing RBCs with diverse morphologies.

**Table 6:** Detection-level evaluation (IoU threshold = 0.5, 100 test images, 6,664 ground-truth boxes).

| Method | Pred | TP | FP | FN | Precision | Recall | F1 |
| --- | --- | --- | --- | --- | --- | --- | --- |
| YOLO26n + DenseNet121 | 6,384 | 6,303 | 81 | 361 | 0.9873 | 0.9458 | 0.9661 |

Table 7 compares the performance of the classification of five phenotypes of sickle RBCs using YOLO26n and a two-step framework where YOLO26n’s built-in classification head is replaced with the proposed DenseNet121 ensemble. In microscopy images of sickle RBCs, dense cell arrangements, small target sizes, and severe class imbalance (DO: ∼70%; E: 1.4%; R: 2.2%) inherently challenge single-step detectors. As shown in Table 7, YOLO26n-only already struggles on minority classes (E: F1,=,0.7732; R: F1,=,0.6186), whereas decoupling detection from classification via a DenseNet121 ensemble operating on individual cropped patches substantially mitigates these limitations. The majority class DO (4,421 samples) already achieves strong performance with YOLO26n-only (F1,=,0.9432), yet the two-step approach further enhances it to 0.9841, indicating refined discrimination even for well-represented categories. More importantly, the improvements are dramatic for minority classes: the E class (91 samples) increases from 0.7732 to 0.9355, the R class (136 samples) from 0.6186 to 0.8789, and the G class (646 samples) from 0.6596 to 0.9370, with substantial gains in both precision and recall. These improvements indicate a significant reduction in both false positives and false negatives for underrepresented categories. The macro-average F1-score increases from 0.7696 to 0.9371, overall accuracy improves from 89.07% to 97.06% (+7.99%), and the weighted-average F1-score rises from 0.8903 to 0.9708, collectively confirming that performance gains are consistent across all classes rather than driven solely by the dominant category. These results strongly support the effectiveness of the two-step framework in enhancing classification accuracy, particularly for minority classes, by leveraging DenseNet121’s ability to extract fine-grained morphological features from individual cell patches.

**Table 7:** Per-class classification performance of the YOLO26n-only baseline versus the proposed two-step framework (YOLO26n + DenseNet121) on 100 full-scope test images. YOLO-only evaluates 6,303 matched boxes via the detector’s own classification head; YOLO26n + DenseNet121 evaluates 6,303 boxes using a 5-fold DenseNet121 ensemble on cropped cell patches. ΔF1 denotes absolute F1 improvement over the baseline. Best results per class are in **bold**.

| Class | Method | Precision | Recall | F1-Score | Samples | ΔF1 |
| --- | --- | --- | --- | --- | --- | --- |
| DO | YOLO26n-only | 0.9351 | 0.9514 | 0.9432 | 4,421 | +0.0409 |
|  | YOLO26n + DenseNet121 | **0.9873** | **0.9810** | **0.9841** | 4,421 |  |
| E | YOLO26n-only | 0.7282 | 0.8242 | 0.7732 | 91 | +0.1623 |
|  | YOLO26n + DenseNet121 | **0.9158** | **0.9560** | **0.9355** | 91 |  |
| ES | YOLO26n-only | 0.8902 | 0.8196 | 0.8535 | 1,009 | +0.0964 |
|  | YOLO26n + DenseNet121 | **0.9513** | **0.9485** | **0.9499** | 1,009 |  |
| G | YOLO26n-only | 0.6997 | 0.6238 | 0.6596 | 646 | +0.2774 |
|  | YOLO26n + DenseNet121 | **0.9299** | **0.9443** | **0.9370** | 646 |  |
| R | YOLO26n-only | 0.5228 | 0.7574 | 0.6186 | 136 | +0.2603 |
|  | YOLO26n + DenseNet121 | **0.8301** | **0.9338** | **0.8789** | 136 |  |
| Macro Avg | YOLO26n-only | 0.7552 | 0.7953 | 0.7696 | 6,303 | +0.1675 |
|  | YOLO26n + DenseNet121 | **0.9229** | **0.9527** | **0.9371** | 6,303 |  |
| Weighted Avg | YOLO26n-only | 0.8919 | 0.8907 | 0.8903 | 6,303 | +0.0805 |
|  | YOLO26n + DenseNet121 | **0.9712** | **0.9706** | **0.9708** | 6,303 |  |
| Accuracy | YOLO26n-only |  | 0.8907 |  | 6,303 | +0.0799 |
|  | YOLO26n + DenseNet121 |  | **0.9706** |  | 6,303 |  |

As shown by one of the images in the test dataset in Fig. 7, both methods yield identical detection performance on this image (Det P = 1.00, Det R = 0.95, TP = 78, FP = 0), with the same four undetected cells (FN1–FN4) located at the image boundary or under occlusion, independent of the classification stage. The key difference lies in morphological classification: the single-stage Yolo26n baseline (A) exhibits more red bounding boxes (Cls Acc = 0.86), indicating misclassifications among morphologically similar classes such as DO, R, and G; whereas the proposed two-stage pipeline (B) produces markedly fewer red boxes (Cls Acc = 0.95), confirming that the dedicated DenseNet121 ensemble effectively resolves these morphological ambiguities. This improvement is consistent across the full validation set, where the two-stage pipeline raises weighted classification accuracy from 89.1% to 97.1%, with F1-scores for the minority classes R and G improving from 0.619 to 0.879 and from 0.660 to 0.937, respectively.

**Figure 7:**
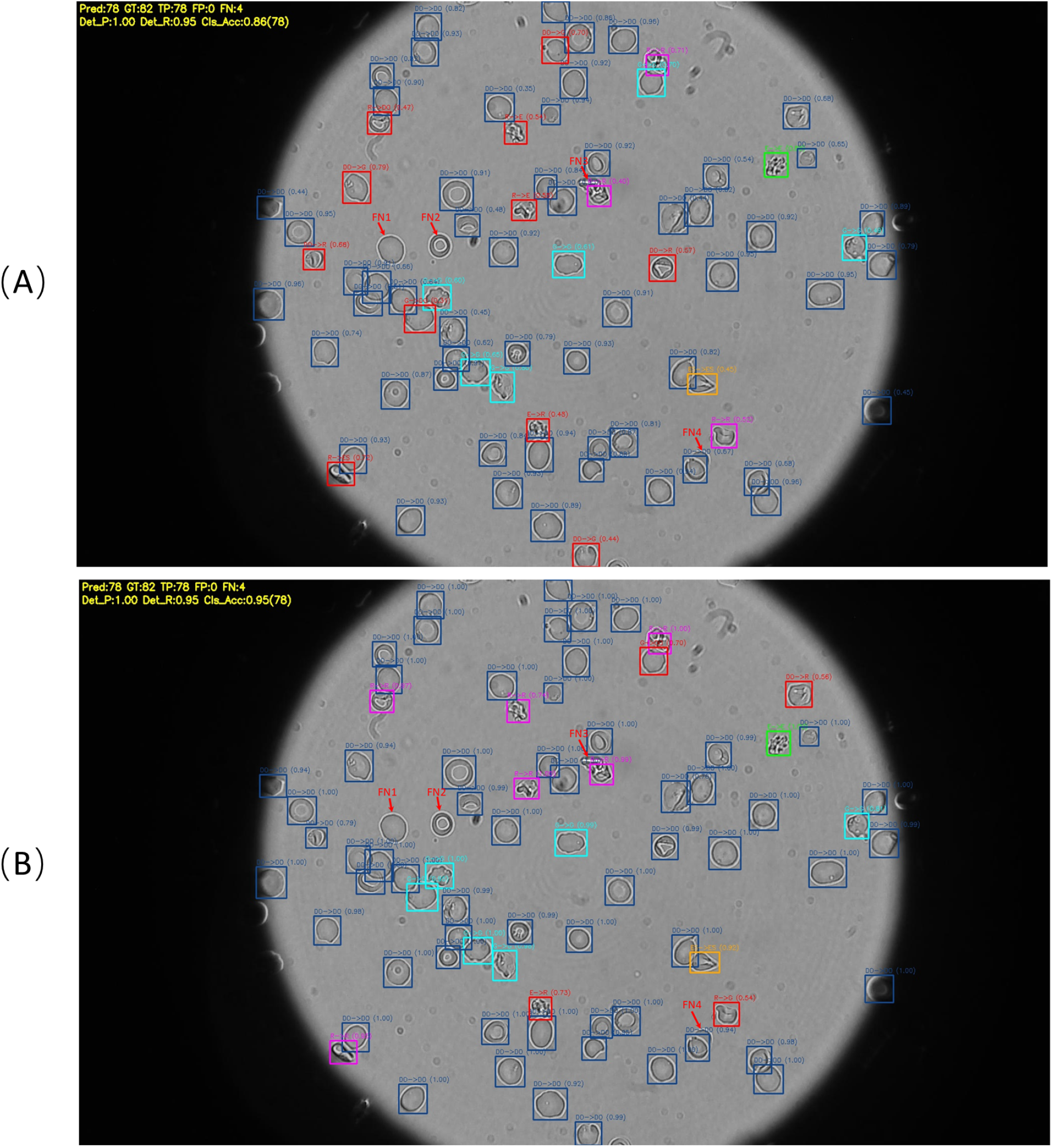
Qualitative performance comparison on a representative brightfield microscopy image beyween **(A)** Single-stage Yolo26n baseline (Cls Acc = 0.86) and **(B)** Proposed two-stage pipeline: Yolo26n + DenseNet121 ensemble (Cls Acc = 0.95). Both methods share identical detection performance (Det P = 1.00, Det R = 0.95); the four undetected cells (FN1–FN4, marked by red arrows) are the same across both methods and correspond to cells at the image boundary or with severe occlusion. Bounding box colors denote the predicted cell morphology: 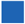 DO, 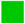 E, 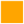 ES, 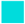 G, 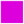 R. 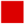 Red boxes indicate misclassified cells (label format: “GT → Pred(Confidence Score)”) or false positives. Comparison between (A) and (B) shows the improved morphological discrimination of the pre-trained DenseNet121 classifier for ambiguous minority classes (G, R, E).

## 4 Discussion and Summary

Our results reveal a consistent and reproducible performance gap between joint detection-classification models and specialized single-cell classifiers when applied to full-scope microscopy images of sickle RBCs. Across 11 model variants from the YOLO and DETR families, strong localization performance did not translate into reliable phenotype classification, particularly for minority classes. Even the best-performing joint model, RF-DETR Small, achieved an mAP@50 of only 0.738 for reticulocytes (R) and 0.882 for granular cells (G), substantially below the classification F1-scores of 0.9333 and 0.9676 achieved by the DenseNet121 classifier on the same phenotypes. This gap persisted across all five imbalance-mitigation strategies we evaluated, with none producing notable gains for the most challenging minority classes. These findings suggest that the performance bottleneck in joint architectures is not primarily a data-quantity problem but reflects a deeper architectural constraint.

This constraint can be understood in terms of task-objective alignment. In joint detection-classification models, the backbone learns feature representations that are shared across both the localization and classification heads. Effective localization requires features that are invariant to cell position, scale, and local density, properties shaped by the bounding box regression loss that dominates training. These same features are then passed to the classification head, which must discriminate among phenotypes based on subtle differences in surface texture, membrane irregularity, and boundary curvature. The problem is that position-invariant features are poorly suited for capturing these morphological details: the spicular projections that distinguish echinocytes, the reticular staining patterns of reticulocytes, and the granular surface texture of G-class cells, all of which require spatially precise representations that detection-oriented models fail to provide. In the two-step framework, this conflict is removed by combining two types of models that can complement each other. The detector is trained as a class-agnostic localizer, enabling it to optimize for recall and box quality without competing classification objectives. The classifier then operates on standardized single-cell patches in which positional variation has been eliminated by cropping, allowing the network to allocate its full representational capacity to morphological discrimination. The dense connectivity of DenseNet121 further supports this by enabling direct gradient flow from the classification loss to early feature layers, facilitating the learning of low-level textural discriminators that joint architectures tend to suppress.

The misclassification patterns in Fig. 6(A) provide additional insight into the remaining classification challenges. The most frequent errors involve confusion between DO and ES (21 instances) and between DO and G (8 instances), suggesting that the classifier sometimes over-weights elongation features when distinguishing normal biconcave morphology from elongated or granular forms. Since DO cells can adopt elongated profiles under certain imaging orientations, these errors likely arise from boundary ambiguity rather than systematic feature confusion. The confusion among E, G, and R classes is more essential: all three exhibit surface texture irregularities that differ in extent, making them inherently harder to separate from morphological features alone. Class E consistently yields the lowest F1-scores across all tested models, which is consistent with its relatively small training set (264 samples after quality control) and its visual similarity to granular cells. Expanding the training data for these minority phenotypes is likely to be the most direct path to further improvement.

In the context of sickle RBC analysis, most existing classification approaches [48–52] report performance on pre-cropped, single-cell images, making direct comparison with our full-scope pipeline non-trivial. However, our classifier component, evaluated under five-fold cross-validation on single-cell crops, achieves accuracy and F1-scores that are competitive with or exceed those reported in prior work, while the full two-step pipeline additionally handles the detection and localization problem that those methods do not address. The practical significance of this is substantial: a model that requires pre-cropped inputs cannot be deployed on raw microscopy data without a separate, manually tuned segmentation step, whereas our framework processes full-scope images end-to-end with inference times below 10 ms per image.

Several limitations of the current work should be acknowledged. First, the dataset originates from a single imaging modality and two institutions, and the generalizability of the trained models to other microscope configurations, staining protocols, or patient populations remains to be established. Second, the classifier was trained and evaluated on static images acquired under normoxic conditions; its performance under dynamic conditions, where RBC morphology changes continuously during deoxygenation and reoxygenation, has not been assessed. Extending the framework to time-lapse data and validating it across multi-center cohorts are important priorities for future work. Third, the current evaluation does not include the clinically relevant task of tracking individual cells across frames, which will require integrating the detection output with a tracking module.

In summary, this work identifies a fundamental limitation of joint detection-classification architectures for fine-grained biomedical image analysis and demonstrates that decoupling the two tasks into separate, specialized components effectively resolves it. The proposed two-step pipeline achieves 97.06% overall classification accuracy and a macro-average F1-score of 0.9371 on full-scope microscopy images of sickle RBCs, with particularly large improvements for minority phenotypes that are clinically relevant to disease prognosis and treatment evaluation. We hope these results and the accompanying benchmark encourage future work on realistic, full-scope evaluation protocols for biomedical cell analysis.

## Acknowledgments

H.L. would like to thank Shan-Ho Tsai from the Georgia Advanced Computing Resource Center (GACRC) for providing technical support.

